# The CCT chaperonin and actin modulate the ER and RNA-binding protein condensation during oogenesis to maintain translational repression of maternal mRNA and oocyte quality

**DOI:** 10.1101/2024.07.01.601596

**Authors:** Mohamed T. Elaswad, Mingze Gao, Victoria E. Tice, Cora G. Bright, Grace M. Thomas, Chloe Munderloh, Nicholas J. Trombley, Christya N. Haddad, Ulysses G. Johnson, Ashley N. Cichon, Jennifer A. Schisa

## Abstract

The regulation of maternal mRNAs is essential for proper oogenesis, the production of viable gametes, and to avoid birth defects and infertility. Many oogenic RNA-binding proteins have been identified with roles in mRNA metabolism, some of which localize to dynamic ribonucleoprotein granules and others that appear dispersed. Here, we use a combination of *in vitro* condensation assays and the *in vivo C. elegans* oogenesis model to determine the intrinsic properties of the conserved KH-domain MEX-3 protein and to identify novel regulators of MEX-3 and the Lsm protein, CAR-1. We demonstrate that MEX-3 undergoes liquid-liquid phase separation and appears to have intrinsic gel-like properties *in vitro*. We also identify novel roles for the CCT chaperonin and actin in preventing ectopic RNA-binding protein condensates in maturing oocytes that appear to be independent of MEX-3 folding. CCT and actin also oppose the expansion of ER sheets that may promote ectopic condensation of RNA-binding proteins that are associated with de-repression of maternal mRNA. This regulatory network is essential to preserve oocyte quality, prevent infertility, and may have implications for understanding the role of hMex3 phase transitions in cancer.

**Significance statement:**

- The molecular mechanisms that regulate phase transitions of oogenic RNA-binding proteins are critical to elucidate but are not fully understood.
- We identify novel regulators of RNA-binding protein phase transitions in maturing oocytes that are required to maintain translational repression of maternal mRNAs and oocyte quality.
- This study is the first to elucidate a regulatory network involving the CCT chaperonin, actin, and the ER for phase transitions of RNA-binding proteins during oogenesis. Our findings for the conserved MEX-3 protein may also be applicable to better understanding the role of hMex3 phase transitions in cancer.

## Introduction

Liquid-liquid phase separation (LLPS) is now understood as a key organizing principle of many cell types, including germ cells. Well over a century ago, the existence of distinct phases in starfish eggs was suggested based on rudimentary imaging (Wilson, 1899); however, only more recently has this been proven. In a pioneering study, germ granules in *C. elegans* were the first RNA granule revealed to be a liquid droplet formed by LLPS (Brangwynne et al., 2009). Numerous studies have now elucidated the essential nature of regulating phase transitions for cellular homeostasis and preventing disease (reviewed in Boeynaems et al., 2023; Phillips and Updike, 2022; Alberti and Hyman, 2021). In some contexts, condensates form aberrantly and are deleterious, e.g. FUS aggregates in ALS patients (Patel et al., 2015), while in other contexts condensates are required for cellular homeostasis, e.g. *osk* RNP complexes in *Drosophila* oocytes (Bose et al., 2022), and yet other examples suggest some condensates are incidental without essential function (Thomas et al., 2023).

During oocyte growth of GV-stage mammalian oocytes, several RNA-binding proteins and maternal mRNAs undergo phase separation to comprise the MARDO (mitochondria-associated ribonucleoprotein domain) (Cheng et al., 2022). When mouse oocytes resume meiosis after the prophase I arrest, the MARDO condensates dissolve. Interestingly, phase transitions of at least two of the MARDO RNA-binding proteins, DDX6 and Lsm14, are described in oocytes of other vertebrate and invertebrate species (Ladomery et al., 1997; Nakamura et al., 2001; Navarro et al., 2001; Audhya et al., 2005; Boag et al., 2005; Squirrel et al., 2006; Jud et al., 2008; Noble et al., 2008; Hubstenberger et al., 2013; Elaswad et al., 2022). In *C. elegans* maturing oocytes DDX6/CGH-1 and Lsm14/ CAR-1 are relatively diffuse and enriched in small granules dispersed throughout the cytosol (Navarro et al, 2001; Boag et al., 2005). However, both proteins undergo dramatic condensation into large RNP granules in arrested oocytes of middle-aged hermaphrodites or female worms lacking sperm (Schisa et al., 2001; Jud et al., 2007; Jud et al., 2008; Noble et al., 2008; Wood et al., 2016). The large RNP granules also include untranslated maternal mRNAs, germ granule proteins (e.g. PGL-1, GLH-1/Vasa), additional Processing-body proteins (e.g. DCAP-1), stress granule proteins (e.g. PABP), and other RNA-binding proteins that are normally diffuse in active oocytes (e.g. MEX-3, LIN-41, OMA-1) (Schisa et al., 2001; Spike et al., 2014 (1513); Tsukamoto et al., 2017). MEX-3 is a conserved KH-domain protein that physically interacts with LIN-41 and OMA-1 (Spike et al., 2014 (1513); Tsukamoto et al., 2017). Its sequence includes multiple intrinsically disordered regions (IDRs) which can drive weak multivalent interactions and LLPS (Elbaum-Garkinkle et al., 2015; Nott et al., 2015; Wood et al., 2016). In addition, two of the four human Mex3 orthologs colocalize with P-body proteins in cell culture, and dysregulated condensation of hMex3A leads to mRNA degradation in colorectal cells and poor survival outcomes for patients (Buchet-Poyeau et al., 2007; Chen et al., 2024). Taken together, MEX-3 phase transitions are conserved and regulated.

Genetic screens have identified regulators of RNA-binding protein condensation, including the chaperonin-containing TCP-1 (CCT/TriC) chaperonin as a promoter of RNA-binding protein condensation in arrested *C. elegans* oocytes (Hubstenberger et al., 2015; Wood et al., 2016). However, no studies to date have identified regulators required to maintain the decondensed state of RNA-binding proteins in the maturing oocytes of young *C. elegans* hermaphrodites. Using a combination of *in vitro* condensation assays and *C. elegans* oogenesis as an *in vivo* model, here we characterize the intrinsic properties of the MEX-3 protein and identify regulators of MEX-3 condensation. MEX-3 undergoes liquid-liquid phase separation similar to the germ granule proteins PGL-1 and MEG-3; however, in contrast to the liquid-like phase of PGL-1, MEX-3 appears to have intrinsic gel-like properties like MEG-3 (Wang et al., 2014; Putnam et al., 2019). We demonstrate a novel role for the CCT chaperonin and actin in preventing ectopic condensates in maturing oocytes that appears to be independent of regulation of MEX-3 folding. The CCT chaperonin and actin concomitantly prevent ectopic ER sheets which may promote the condensation of RNA-binding proteins by restricting molecular diffusion to a two-dimensional plane and reducing the concentration threshold for condensation. Lastly, our results suggest inhibiting ectopic RNA-binding protein condensates and ER sheets in maturing oocytes may be essential to maintain the repression of maternal mRNA translation and preserve oocyte quality.

## Results

### MEX-3 undergoes liquid-liquid phase separation and behaves as a gel-like phase in vitro

To better understand the physicochemical basis for the dynamics of MEX-3 in the germline and early embryo *in vivo*, we asked if recombinant MEX-3 undergoes LLPS *in vitro.* Based on the intrinsically disordered regions (IDRs) predicted by AlphaFold-2 and the MobiDB database, we predicted that MEX-3 undergoes LLPS (Figure 1A; Piovesan et al., 2022; Piovesan et al., 2023). We purified recombinant MEX-3 and fluorescently labeled it using DyLight-488 (Supplementary Figure 1, A and B). We first collected images of MEX-3 in condensation buffer at a range of concentrations and a range of salt concentrations. Droplets spontaneously increased in size and number as the protein concentration increased and the salt concentration decreased (Figure 1, B and C; Supplementary Figure 1C). At the physiological MEX-3 concentration in oocytes, estimated to be 0.1 µM, we detected rare, small droplets (Figure 1C). This concentration was extrapolated from the MEX-3 concentration in early embryo extracts of 0.173 µM (Saha et al., 2016). Our observations suggest MEX-3 *in vivo* is poised to undergo phase separations. Including and increasing the concentration of the crowding agent PEG8000 resulted in increased turbidity that was visible in the tubes and greater numbers of condensates (Figure 1D; Supplementary Figure 1D). The inclusion of 10% PEG with 1.0 µM MEX-3 resulted in large, non-spherical, fibrillar structures (Figure 1D). Taken together, the increase in MEX-3 droplet size in response to changes in salt and protein concentrations, and to the presence of a crowding agent, suggest that MEX-3 undergoes liquid-liquid phase separation.

**Figure 1.**
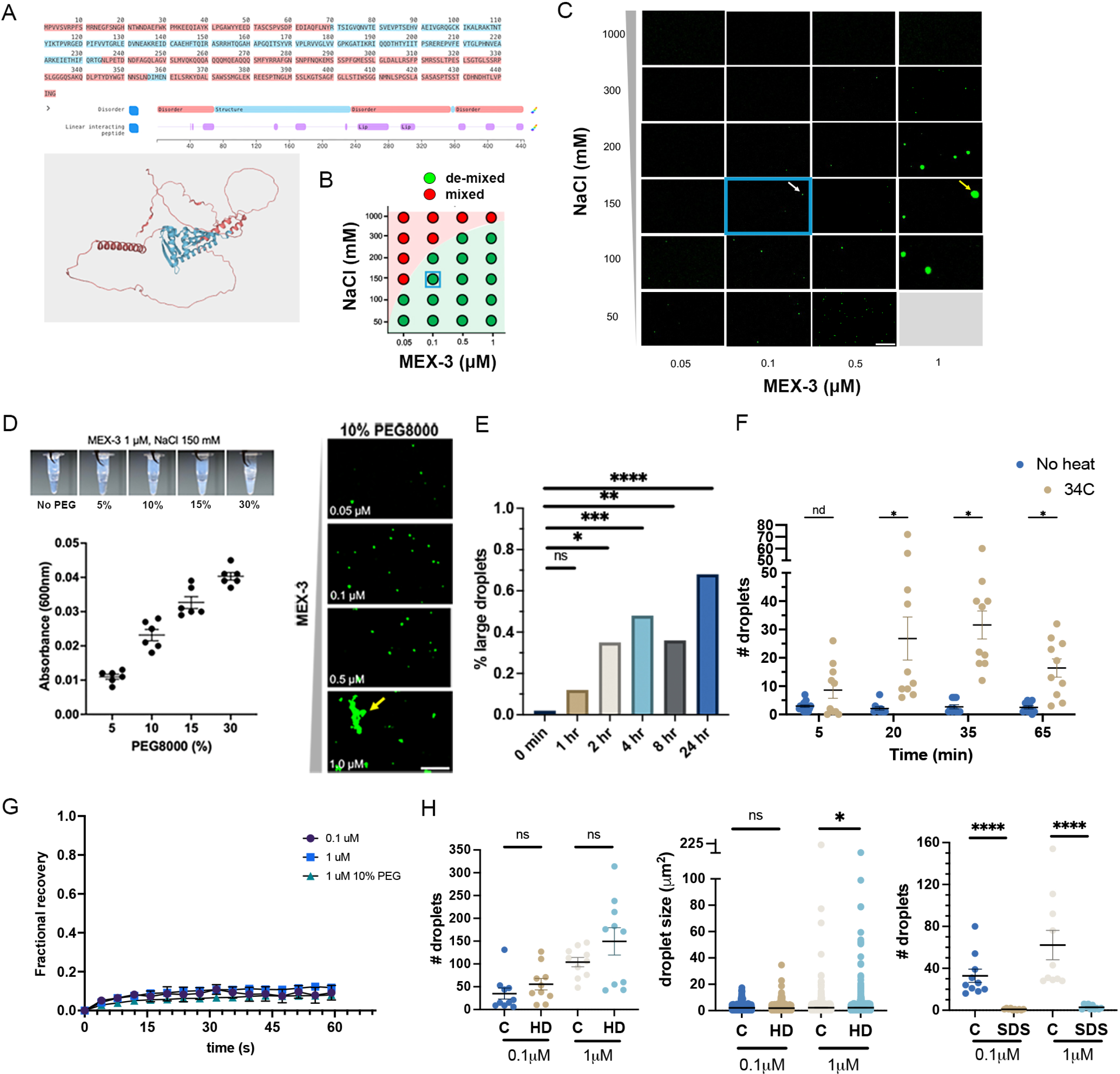
MEX-3 undergoes liquid-liquid phase separation and behaves as a gel-like phase. A) The MobiDB tool identifies MEX-3 as having multiple linear interacting peptides and intrinsically disordered regions (IDRs) which are seen as unstructured loops in the AlphaFold2 image. B) Phase diagram shows threshold between mixed states with no phase separation (red), and de-mixed states with phase separation (green). C) Fluorescently-labeled, recombinant MEX-3 imaged in buffers at a variety of protein and salt concentrations. White arrow in blue box indicates small droplet at physiological salt and MEX-3 concentrations. Yellow arrow indicates large droplet at high MEX-3 concentration of 1 µM. D) *Left* Addition of PEG8000 at increasing concentrations results in visibly turbid MEX-3 in buffer; absorbance measured at 600 nm. *Right* Increasing concentrations of MEX-3 in the presence of 10% PEG results in large fibrillar structures (yellow arrow). E) The % of droplets of 0.1 µM MEX-3 larger than 20.3 µm^2^ increases over time. F) The # of droplets of 0.1 µM MEX-3 was determined within 5-, 20-, 35-, and 65-min exposure to 34°C and compared to no-heat controls imaged in parallel. G) Fractional recovery after FRAP of MEX-3 droplets at three concentrations indicates immobile protein. H) MEX-3 droplets at 0.1 µM and 1 µM in the presence of 10% PEG were counted after exposure to 10% hexanediol or 10% SDS detergent. Scale bar is 20 µm in all micrographs. In panels E and H, error bars are mean +/-SEM. ns is not significant; * *p*<.05; \*\*\**p*<0.001; \*\*\*\**p*<0.0001 by Mann-Whitney or Kruskal-Wallis test. In panel F, ns is no discovery; * is a discovery by Mann-Whitney tests and the False Discovery Rate-two stage step up test. Note disrupted Y-axis in panels F and H (middle).

To begin to assess if purified MEX-3 droplets are liquid-like, we first imaged 0.1 µM MEX-3 after 15 or 30 min in buffer. The small, spherical MEX-3 droplets appeared stable once we began imaging (within ∼1 min), and we did not detect any fusion events within the first few mins that would be consistent with a liquid phase growing by Ostwald ripening and fusion. We also did not detect large changes in droplet size or shape after 15 or 30 mins (Supplementary Figure 2, A and B; therefore, we conclude MEX-3 does not behave as a liquid phase *in vitro.* We next asked if MEX-3 droplets change in size or shape over a time-scale of hours. We detected a significant increase in droplet size and an increase in non-spherical droplets starting at 2 hours, both of which suggest MEX-3 is somewhat dynamic with gel-like properties *in vitro* (Figure 1E; Supplementary Figure 2C). Secondly, we asked if MEX-3 droplets exhibit the sensitivity to high temperatures typically seen for proteins in liquid-like phases (Putnam et al., 2019; Watkins and Schisa, 2021). We imaged 0.1 µM MEX-3 after 15-60 min of heat exposure and quantitated the number of condensates compared to controls. Instead of dissolving, the number of MEX-3 droplets significantly increased within 15 min of exposure to elevated temperature (Figure 1F). Thirdly, to determine the mobility of MEX-3 *in vitro*, we performed FRAP analyses of small droplets at 0.1 µM, larger droplets at 1.0 µM, and fibrillar structures at 1.0 µM + 10% PEG. In all three conditions, MEX-3 failed to recover, suggesting a non-dynamic phase and consistent with a gel-like or solid phase (Figure 1G). Lastly, we asked if exposure of 0.1 µM or 1 µM MEX-3 droplets to the aliphatic alcohol 1,6-hexanediol (HD) or SDS detergent was sufficient to dissolve droplets. We performed assays in buffers with 10% PEG to start with sufficient numbers of droplets. At both MEX-3 concentrations, the numbers of droplets increased slightly but not significantly after HD exposure, and droplet size decreased slightly using 1 µM MEX-3 (Figure 1H). The mild sensitivity to HD suggests the droplets are not liquid phases. In contrast, after exposure to the detergent SDS droplets were nearly undetectable, indicating the droplets are not solid phases (Figure 1H). Collectively, the results from these four assays are most consistent with MEX-3 having intrinsic gel-like properties *in vitro*.

### The CCT chaperonin prevents ectopic condensation of MEX-3 in maturing oocytes

We next investigated the regulation of MEX-3 *in vivo*. To follow up on the identification of the CCT chaperonin as a regulator of RNA-binding proteins in *C. elegans* arrested oocytes and embryos, we asked if CCT regulates the phase of MEX-3 protein in maturing oocytes by performing RNAi by feeding in the *gfp*-tagged *mex-3* allele (Tsukamoto et al., 2017). After depletion of *cct-1*, we detected heterogeneous clusters of GFP::MEX-3 protein, in contrast to the diffuse MEX-3 in control oocytes (Figure 2A). Clusters were often cortically enriched or associated with the nuclear envelope, and some contained small, spherical puncta. We also noted variably altered oocyte morphology, likely due to the critical role of CCT in folding actin and tubulin monomers and regulating the cell cycle (Frydman et al., 1992; Vinh and Drubin, 1994; Camasses et al., 2003). We were unable to examine oocytes in a mutant *cct* allele, as existing null alleles are larval lethal (Wu and Herman, 2006; C. elegans Knockout Consortium). To characterize the heterogeneous shapes of clusters, we measured their circularity after *cct-1* depletion and found a moderate, but significant negative correlation of 0.6691 between circularity and size (Spearman, p<0.0001; Supplementary Figure 3A), indicating the largest clusters tend to be less circular than small clusters. This result suggests CCT-1 promotes a diffuse state of MEX-3 in maturing oocytes, opposite from its role promoting condensed phases of proteins in arrested oocytes and early embryos, but similar to the role of CCT inhibiting ectopic P-bodies and polyQ aggregates (Tam et al., 2006; Nadler-Holly et al., 2012; Updike and Strome, 2009; Hubstenberger et al., 2015; Wood et al., 2016).

**Figure 2.**
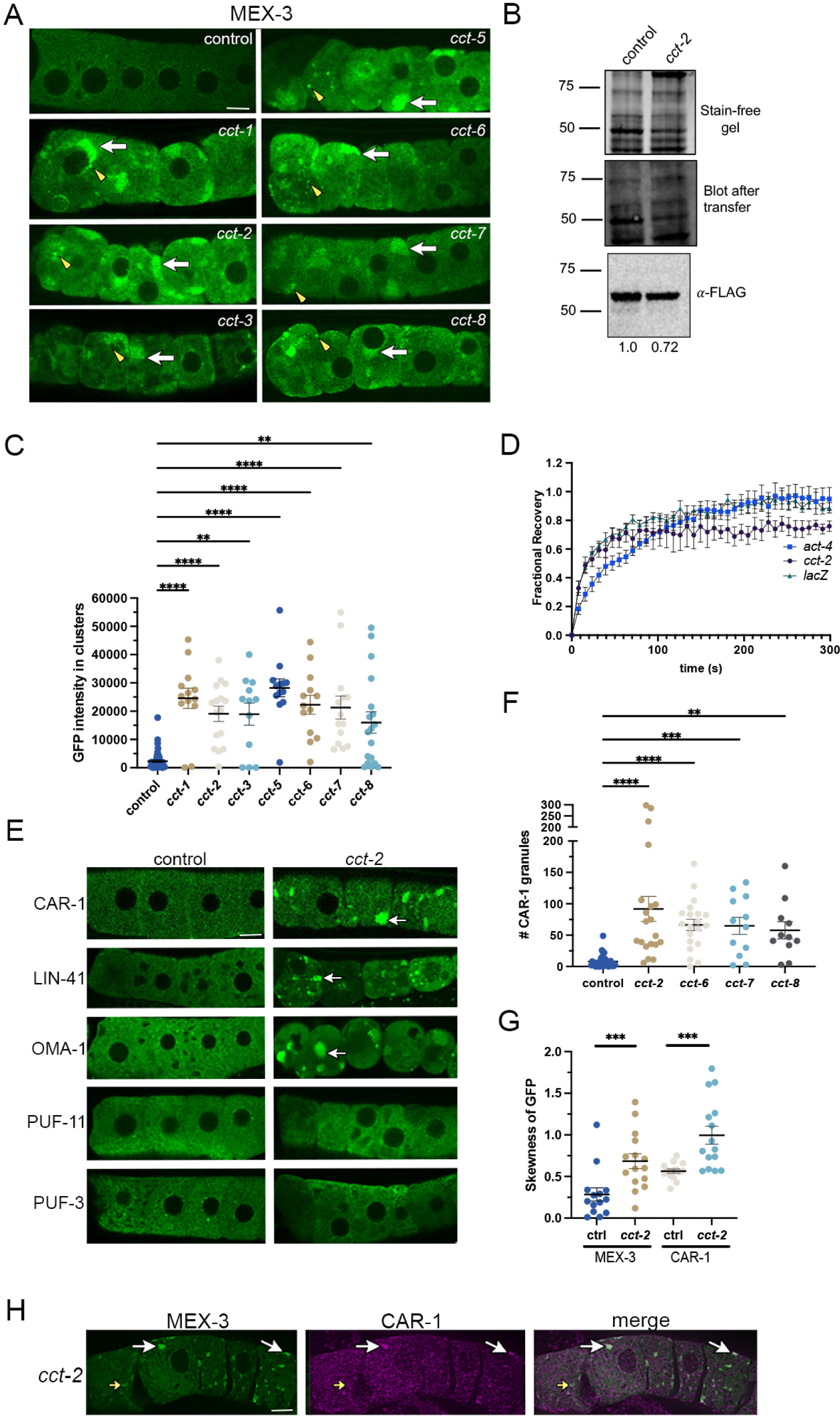
The CCT chaperonin inhibits ectopic condensation of MEX-3 in maturing oocytes. A) Confocal single slices showing GFP::MEX-3 in proximal oocytes after depletion of individual *cct* subunits by RNAi compared to the control of *lacZ(RNAi)*. Here, and in all micrographs, the most proximal oocyte is oriented to the left. White arrows indicate large protein clusters; yellow arrowheads indicate small, spherical clusters. In all images of oocytes, a mid-focal Z-slice is shown where nuclei of only a subset of oocytes may be visible. B) Representative Western blot of CCT-6::3xFLAG worm extracts treated with *lacZ* control RNAi or *cct-2(RNAi)*, 50 worms/lane. Band density was quantified using Bio-Rad Image Lab5.1 software. The intensity of the FLAG-tagged CCT-6 was normalized to total protein transferred to the blot, and the intensity relative to the control is shown under each lane. Molecular weight is shown to the left in kD. C) Significant increases in GFP::MEX-3 intensity in clusters after RNAi depletion of *cct* subunits was demonstrated by particle analysis. D) Fractional recovery after FRAP to assess mobility of GFP::MEX-3 after RNAi of *act-4, cct-2,* or *lacZ* control. E) Confocal single slices showing GFP-tagged RNA-binding proteins after depletion of *cct-2* compared to the *lacZ* control. White arrows indicate large protein clusters. F) Quantitation of numbers of CAR-1 granules after depletion of four *cct* subunits. Note y-axis is disrupted. G) The Fiji skewness tool measured the average uniformity of MEX-3::GFP and CAR-1::GFP within the three most-proximal oocytes after depletion of *cct-2* and *lacZ* and revealed similar significant differences as particle analyses. H) Immunostaining for CAR-1 (magenta) in GFP::MEX-3 strain (green) after depletion of *cct-2* by RNAi. White arrows indicate condensates. Scale bar is 10 µm in all micrographs. In all graphs, error bars are mean +/-SEM. \*\**p*<0.01; \*\*\**p*<0.001; \*\*\*\**p*<0.0001, by Mann-Whitney or Kruskal-Wallis tests.

The CCT chaperonin comprises eight subunits that share ∼30% sequence identity (Kubota et al., 1994, 1995). It was not obvious if all eight CCT subunits would be required to regulate MEX-3 in oocytes for two reasons. First, distinct subsets of CCT chaperonin subunits were previously identified as regulators of MEX-3, CAR-1, and PGL-1, suggesting specific subunits of the CCT chaperonin may modulate each RNA-binding protein as shown for CCT-3 (Nadler-Holly et al., 2012). Second, CCT subunits can function as monomers in addition to their folding activity as part of the CCT chaperonin (Elliot et al., 2015; Spiess et al., 2015; Ma et al., 2022). We therefore decided to deplete CCT subunits individually to try to determine if CCT chaperonin oligomer activity or CCT-1 specifically prevents ectopic clusters of MEX-3 in oocytes. In siRNA studies, targeting of single CCT subunits results in variably decreased levels of non-targeted CCT subunits (Brackley and Grantham, 2010; Elliot et al., 2015). In addition, off-target silencing of closely related sequences can occur during *C. elegans* RNAi, and the eight *cct* sequences are 40-50% identical to one another (Fire et al., 1998; Rual et al., 2007).. Thus, we first asked if RNAi of a single CCT subunit affects the levels of other CCT subunits. We created a FLAG-tagged allele of *cct-6*, and after RNAi of *cct-2*, our semi-quantitative Western analysis revealed levels of CCT-6 reduced to 0.72 of the levels in the control (Figure 2B). This result suggests levels of non-targeted *cct* subunits are modestly depleted in our RNAi system. After depleting six additional *cct* subunits individually, we detected heterogeneous MEX-3 cluster sizes and shapes, similar to *cct-1* RNAi (Figure 2B). We determined the average fluorescence intensity of MEX-3 in clusters was significantly increased after depletion of each subunit (Figure 2C). We interpret these results as consistent with a model where the folding activity of the CCT chaperonin oligomer is required to prevent ectopic clusters of RNA-binding proteins in maturing oocytes. We confirmed CCT chaperonin folding activity was disrupted after *cct-2* RNAi after detecting a 2-fold reduction in levels of its obligate substrate β-tubulin and decreased amounts of microtubule polymerization in 75% of GFP::TBB-2 gonads (n=16) (Supplementary Figure 3, B and C).

If MEX-3 is folded by CCT, the ectopic clusters of MEX-3 after depletion of *cct* subunits could result from unfolded MEX-3 protein aggregates. Alternatively, MEX-3 may not be a substrate of CCT, and the clusters could be ectopic MEX-3 condensates. MEX-3 undergoes condensation when sperm become depleted in hermaphrodites and meiotic maturation arrests (Schisa et al., 2001; Jud et al., 2008; Elaswad et al., 2022). We performed FRAP analyses after *cct-2* RNAi to characterize the mobility of MEX-3 in ectopic clusters. MEX-3 recovered very quickly after photobleaching, with 68% fractional recovery within 50 seconds, and we did not detect any notable changes from the control experiment, *lacZ* RNAi (Figure 2D). We conclude MEX-3 is normally mobile when in a diffuse state in maturing oocytes and maintains a highly mobile phase in *cct-* induced ectopic clusters. Since unfolded protein aggregates are generally expected to be immobile in FRAP assays, we interpret the results to strongly suggest that ectopic MEX-3 clusters are condensates (Kim et al., 2002; Roberti et al., 2011; Saegusa et al., 2014).

### The CCT chaperonin is required to prevent ectopic condensation of multiple RNA-binding proteins in maturing oocytes

We next asked if the CCT chaperonin selectively inhibits MEX-3 condensation, or if it is required to maintain decondensed phases of additional RNA-binding proteins in maturing oocytes. We first tested the effect of depleting *cct* subunits in GFP::CAR-1 oocytes. CAR-1 has homology to the Sm domain proteins Trailerhitch and Lsm14 and is required for oogenesis (Boag et al., 2005). In control oocytes CAR-1 was largely decondensed with small puncta and modest membrane enrichment as expected. After depletion of *cct-2*, numerous, heterogeneous CAR-1 clusters were detected (Figure 2E; Boag et al., 2005). Similar penetrant CAR-1 phenotypes were observed after depletions of additional *cct* subunits (Figure 2F). We confirmed the disruption of the decondensed state for CAR-1 and MEX-3 after *cct* depletion using skewness analysis (Figure 2G). To determine if MEX-3 and CAR-1 condense in the same RNP complexes, we did immunostaining for CAR-1 in the *gfp*-tagged *mex-*3 allele after *cct-2* depletion, and although we had to use suboptimal fixation conditions for CAR-1 to maintain the GFP signal, we observed several examples where MEX-3 and CAR-1 appear to co-localize (Figure 2H).

We next asked if CCT regulates phase transitions of LIN-41 and OMA-1, two components of ribonucleoprotein particles in oocytes that include MEX-3 (Spike et al., 2014). LIN-41 (a TRIM-NHL protein) and OMA-1 (a TIS11, Zn-finger RNA-binding protein) are required for oogenesis and repress translation of several maternal mRNAs (Detwiler et al., 2001; Spike et al., 2014a; Spike et al., 2014b). After depletion of *cct-2*, both proteins appeared in highly penetrant, ectopic condensates that were not detected in control oocytes (Figure 2E; 72%, n=21 for LIN-41; 87%, n=25 for OMA-1). PUF-3 and PUF-11 are two homologs of Pumilio proteins that contribute to translational repression of similar target mRNAs as LIN-41; however, we did not detect any notable condensates of either PUF-3 or PUF-11 after *cct-2* depletion (Figure 2E; n=18, 26). Because we still observed variable disorganization of the oocytes in these experiments, the PUF-3 and PUF-11 results suggest that ectopic condensation of RNA-binding proteins does not occur solely as a consequence of oocyte disorganization induced by *cct* depletion. We next explored the possibility that *cct-*depleted oocytes might have increased amounts of yolk, where the yolk could act as intracellular crowding agent to induce clusters of RNA-binding proteins. We depleted *cct* in the VIT-2::GFP strain, a reporter for the vitellogenin protein abundant in yolk (Grant and Hirsch, 1999). Not only did we not detect any increase in the amount of VIT-2 in oocytes, we detected high amounts of yolk in the hypodermis outside of the germline and only very low levels of yolk within *cct* oocytes (Supplementary Figure 3D). This result suggests depletion of *cct* blocks transport of yolk from the intestine to the oocyte, perhaps because actin, an obligate substrate of the CCT chaperonin, is required for receptor-mediated endocytosis as it is in organisms from yeast to mammalian cells (Lamaze et al., 1997; Wendland et al., 1998).

Taken together, the CCT chaperonin appears to be required to prevent ectopically condensed phases of a subset of critical RNA-binding proteins in oocytes. Given the FRAP data where MEX-3 remains mobile within ectopic condensates which is largely inconsistent with unfolded protein aggregates, it is possible none of these RNA-binding proteins are substrates of CCT. Instead, the condensation phenotypes may be due to one or more intermediates that require CCT to achieve their native structure and function in modulating phase transitions.

### CCT is expressed at high levels in the germline and functions cell autonomously

In mammalian cells, the CCT chaperonin is broadly expressed throughout the cytosol and is also detected in some nuclei (Joly et al., 1994; Roobol et al., 1999). Prior approaches to determine the expression pattern of *cct* subunits in *C. elegans* relied on promoter fusions that detected expression in muscle, hypodermal, and neuronal cells, but precluded analyses of the germline (Leroux and Candido, 1997). To determine the subcellular distribution of the CCT chaperonin in the germline, we created a FLAG-tagged allele of *cct-6* and did anti-FLAG immunostaining. We detected high levels of CCT-6 in oocytes and throughout the germline (Supplementary Figure 4A). Levels were severely decreased after depletion of *cct-6*, demonstrating the staining was specific to CCT-6. Based on its high levels of expression in the germline, we predicted that the CCT chaperonin functions cell autonomously in the germline to modulate phase transitions. Therefore, we did CAR-1 immunostaining after depleting *cct-2* in an allele of *rde-1* that induces germline-specific RNAi (Zou et al., 2019). RDE-1 encodes an Argonaute that functions cell autonomously in RNAi (Tabara et al., 1999). We detected numerous CAR-1 condensates in *cct-2-*depleted oocytes (Supplementary Figure 4B), suggesting that CCT functions in the germline to regulate phase transitions of oocyte RNA-binding proteins.

### Actin is required to maintain decondensed phases of RNA-binding proteins in oocytes

Proteins can condense into liquid or gel droplets when physico-chemical conditions change (Hyman et al., 2014; Alberti, 2017). Therefore, we asked if condensates caused by depletion of *cct* subunits might result from increased levels of MEX-3. We performed semi-quantitative Western blots and observed a moderate increase in the levels of MEX-3 after *cct* depletion compared to the control (Figure 3A). This result suggests increased protein concentration may contribute to the condensation phenotype. Considering the CCT chaperonin is estimated to fold ∼9-15% of the proteome (Thulasiraman et al., 1999), it also seems possible that one or more of its substrates may contribute to the regulation of phase transitions independent of modulating protein levels.

**Figure 3.**
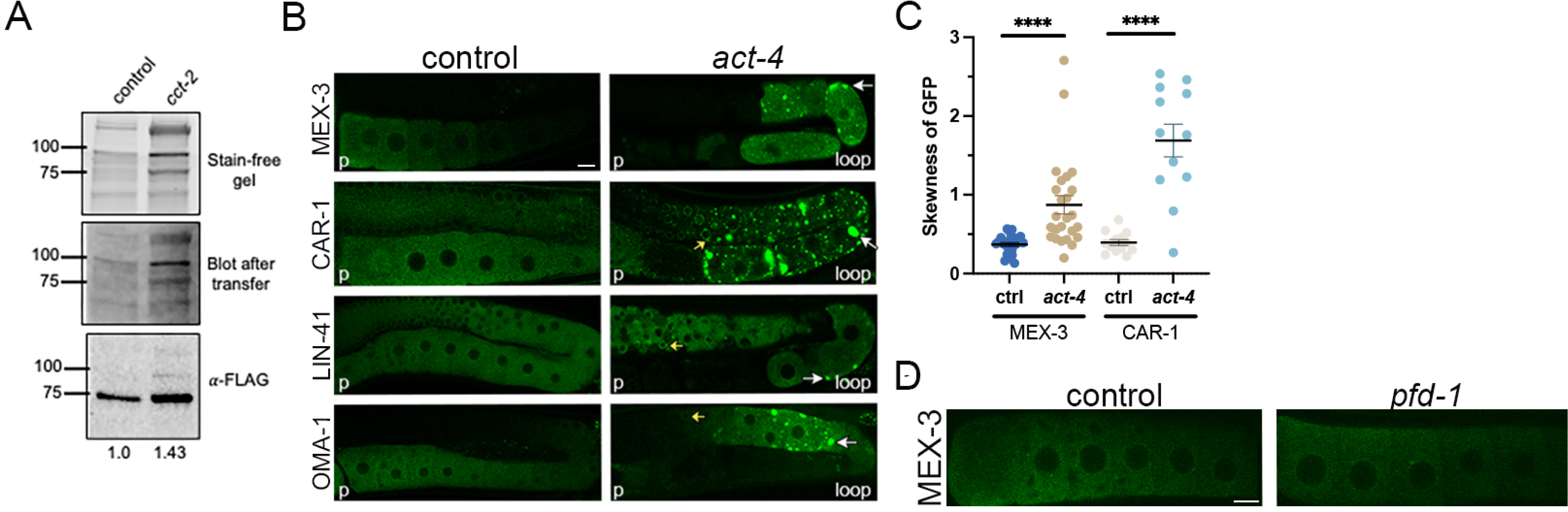
Actin is required to maintain decondensed phases of RNA-binding proteins in oocytes. A) Representative Western blot of 3xFLAG::MEX-3 worm extracts treated with *lacZ* control RNAi or *cct-2(RNAi)*, 50 worms/lane. Band density was quantified using Bio-Rad Image Lab5.1 software. The intensity of the FLAG-tagged MEX-3 was normalized to total protein transferred to the blot, and the intensity relative to the control is shown under each lane. Molecular weight is shown to the left in kD. B) Confocal single slices showing GFP-tagged RNA-binding proteins in germlines after *act-4* or *lacZ* control depletions by RNAi. In controls the most proximal oocyte is labeled p. In *act-4* germlines, the location of the proximal germline (similar to an empty gonad phenotype lacking cellularized oocytes) is labeled p, and the loop with cellularized oocytes is labeled. White arrows indicate condensates. C) Cellularized oocytes at the loop were analyzed after *act-4(RNAi).* Skewness analysis indicates the distribution of MEX-3 and CAR-1 became significantly less uniform throughout the cytosol after *act-4(RNAi)*. Error bars are mean +/-SEM. \*\*\*\**p*<.0001 by Mann-Whitney tests. D) Confocal slices of GFP::MEX-3 in oocytes of control and *pfd-1(RNAi)* worms. Scale bar is 10 µm in all micrographs.

The best-described substrates of CCT include actin and tubulin monomers (Frydman et al., 1992; Gao et al., 1992). Therefore, we decided to begin to test the hypothesis that CCT modulates phase transitions of RNA-binding proteins via the actin cytoskeleton by using RNAi to deplete *act-4* in GFP::MEX-3 and GFP::CAR-1 worms. Given the strong sequence identity among actin paralogs in *C. elegans*, we predict all actin monomers in the gonad are depleted in this experiment (Fire et al., 1998; Rual et al., 2007). Depletion of actin resulted in pleiotropic effects on gonad architecture, including an empty-gonad phenotype in the proximal region where maturing oocytes are usually found, and cellularized oocytes near and at the loop (Velarde et al., 2007; Wolke et al., 2007). We detected ectopic MEX-3 condensates and numerous CAR-1 condensates in the cellularized oocytes and quantitated the degree of condensation to that of the most proximal three oocytes of the control (Figure 3B, C). Some ectopic CAR-1 condensates were quite large, and CAR-1 was also ectopically enriched in perinuclear granules of distal cells (Figure 3B). We conclude the actin cytoskeleton is required to inhibit ectopic condensation of MEX-3 and CAR-1. If actin mediates the effect of CCT on RNA-binding protein condensates, we predicted ectopic condensation of LIN-41 and OMA-1 after *act-4* depletion. We detected a low penetrance phenotype of mild-modest LIN-41 condensation (Figure 3B, 31%; n=19) and highly penetrant OMA-1 condensates (Figure 3B, 75%; n=13). These results are consistent with CCT acting in parallel to and/or through actin to modulate RNA-binding protein condensation.

If the *cct*-induced ectopic condensation phenotype is mediated by actin, we would predict two additional outcomes: 1) the *actin*-induced condensates would have similar mobility as *cct-*induced condensates, and 2) depletion of prefoldin would elicit similar phenotypes as actin depletion. Prefoldin is an obligate co-chaperone of CCT in folding newly synthesized actin and tubulin monomers (Vainberg et al., 1998). We performed FRAP analyses after *act-4* RNAi to characterize the mobility of MEX-3 in ectopic clusters and observed nearly-complete recovery after photobleaching. Similar to our results after *cct* depletion, MEX-3 recovered to ∼50% of pre-bleach levels within 50 seconds after *act* depletion (Figure 2D). We used RNAi to deplete four of six *pfd* subunits individually and examined MEX-3 in oocytes. The depletions appeared to be partially effective as we observed decreased amounts of polymerized microtubules in 20-47% of gonads and only slightly decreased levels of GFP::TBB-2 in oocytes (Supplementary Figure 5, A-C); however, we did not detect MEX-3 condensation phenotypes in any oocytes (Figure 3D). Although we can’t be sure levels of prefoldin were adequately depleted to disrupt folding of actin monomers, taken together, these results suggest CCT may act at least partially independently of actin to inhibit ectopic condensation of RNA-binding proteins in maturing oocytes.

### Expanded sheets of ER may promote condensation of RNA-binding proteins in oocytes

In oocytes arrested in prophase I for an extended time, large RNP condensates are detected that can be up to 20x larger than P granules in maturing oocytes (Schisa et al., 2001; Elaswad, Watkins et al., 2022). At the same time, the endoplasmic reticulum (ER) undergoes dramatic remodeling into large sheets that have been proposed to promote the assembly of large RNP condensates (Patterson et al., 2011; Langerak et al., 2017). We therefore asked if ectopic ER sheets form in oocytes concomitant with ectopic RNA-binding protein condensates after depletion of *cct-2* and *act-4*. To visualize the ER, we used a strain with GFP-tagged SP12, a resident signal peptidase that removes signal peptides from proteins as they exit the ER (Poteryaev et al., 2005). In control oocytes, the majority of the ER appeared uniformly distributed throughout the cytosol of oocytes, with some enrichment cortically (Figure 4A). In stark contrast, after depleting subunits of the CCT chaperonin or actin, ectopic sheets and tubules of ER were detected throughout the oocytes, which were scored in the cellularized oocytes at/near the loop after actin depletion (Figure 4A). The uniform distribution of the ER throughout the cytosol was dramatically disrupted in most *cct-* or *actin-*depleted oocytes (Figure 4B; Supplementary Figure 3E; 82%; n=11 for *cct*; 85%; n=20 for *actin*). The correlation between increased ER sheets and ectopic RNA-binding protein condensation after *cct* depletion suggested the possibility that some of the ER was annulate lamellae, a specialized form of ER containing nuclear pore complexes (Kessel et al., 1989). Previous studies show cytoplasmic foci of nucleoporins (Nup) form during oogenesis (Pitt et al., 2000; Thomas et al., 2023). Models have been suggested where annulate lamellae may recruit RNA-binding proteins to assemble into large condensates during stress (Patterson et al., 2011). To ask if Nup foci increase after *cct* depletion, we used a strain with GFP tagged to the Nup358 homolog *npp-9* (Voronina and Seydoux, 2010). In control oocytes, GFP:NPP-9 was detected strongly at the nuclear envelope and in punctate foci in the cytosol as expected (Figure 4C). After *cct* depletion, NPP-9 was still detected at the nuclear envelope. In contrast to the control, we detected significantly fewer NPP-9 foci and in some oocytes increased NPP-9 levels throughout the cytosol (Figure 4C). Dramatic decreases in Nup foci formation occur when a subset of Nups is depleted (Thomas et al., 2023); therefore, the *cct-*induced phenotype may be caused by a reduction in the concentration of certain Nups below the saturation concentration required for condensation. We conclude the dramatically expanded ER sheets do not contain Nup358 and are unlikely to be annulate lamellae.

**Figure 4.**
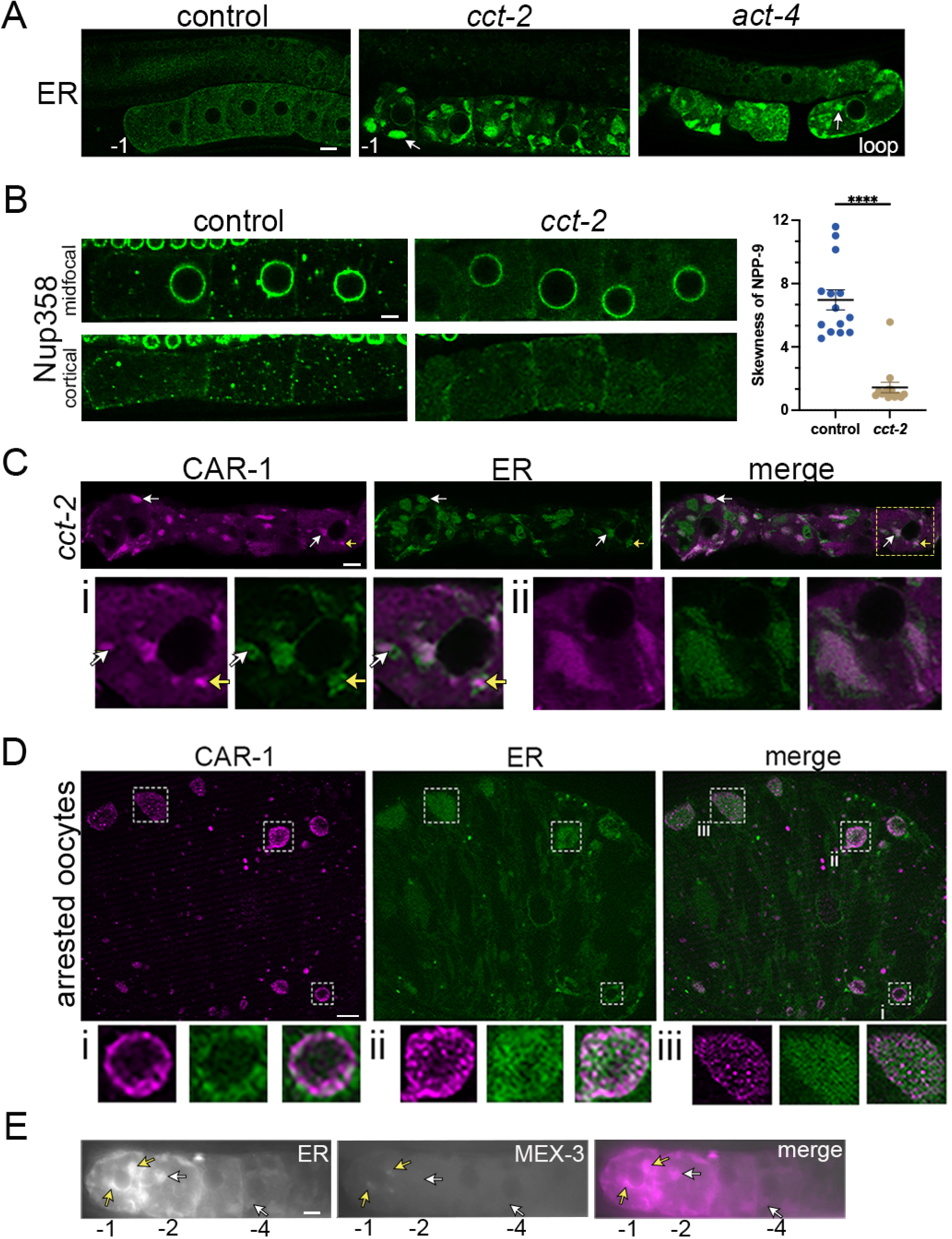
Expanded sheets of ER may promote condensation of RNA-binding proteins in oocytes. A) Confocal slices show GFP::SP12 (ER marker) in oocytes after depletion of *cct-2* and *act-4*. The most proximal oocytes are labeled -1 in the control and *cct-2* image; the loop with cellularized oocytes is labeled in the *act-4* image. White arrows indicate expanded ER sheets. B) Confocal slices show GFP::NPP-9 (Nup358) in oocytes after depletion of *cct-2* or *lacZ* control. Skewness analysis shows significantly fewer NPP-9 foci in mid-focal slices after *cct-2* RNAi. C) Immunostaining of GFP::SP12 oocytes using anti-CAR-1 to visualize condensates after depletion of *cct-2*. Fixation was adapted to allow GFP signal (green is ER) to persist while visualizing CAR-1 (magenta). White arrows indicate CAR-1 condensates adjacent to or partially co-localizing with ER sheets; yellow arrows indicate condensate enclosed by ER. i) Inset of square region at higher magnification; ii) High magnification view of different oocyte from same worm where large CAR-1 condensates appear closely associated with ER sheets. D) Immunostaining using anti-CAR-1 in *fog-2;* GFP::SP12 to assess condensates and ER in arrested oocytes. Insets correspond to labeled regions in merge. i) ER appears to surround some hollow condensates; ii and iii) ER closely associates with condensates. E) Fluorescence micrographs after *cct-2(RNAi)* in oocytes expressing mCherry::SP12 (ER) and GFP::MEX-3. All arrows indicate regions of large ER sheets. In the -1 oocyte, a few ectopic MEX-3 granules (weak *cct* phenotype) correspond to the locations of ER sheets (yellow arrows). In the -2 and -4 oocyte, ER sheets are detected but not MEX-3 granules (white arrows). In all graphs, error bars are mean +/-SEM. \*\*\*\**p*<.0001 by Mann-Whitney tests. Scale bar is 10 µm in all micrographs.

Recent studies have revealed that membrane surfaces can promote condensation, and condensates can modulate membrane structure and function (Snead and Gladfelter, 2019). If ER sheets and RNA-binding protein condensates regulate one another in any manner, we predicted some RNA-binding protein condensates would be in close contact with ER sheets. CAR-1/Trailerhitch colocalizes with the ER in interphase of early embryos, and in nurse cells/oocytes, respectively (Wilhelm et al., 2005; Squirrel et al., 2006). To simultaneously visualize the ER and CAR-1 after depletion of *cct-2*, we performed immunostaining for CAR-1 in GFP:SP12 worms using the same fixation method as in Figure 2H. We noted the majority of CAR-1 condensates, including those at the cortex or nuclear envelope, appear to coincide with ER sheets (Figure 4D). At high magnification, intricately folded ER structures were revealed, some of which appeared to wrap around the smaller CAR-1 condensates and others appeared adjacent to CAR-1 condensates (Figure 4Di). In other oocytes, large CAR-1 condensates appeared to stretch across large ER sheets (Figure 4Dii). We next asked if ER sheets and CAR-1 condensates partially co-localize in a second developmental context where both occur, arrested oocytes of *fog-2* females (Jud et al., 2008; Noble et al., 2008; Langerak et al., 2018). We performed immunostaining for CAR-1 in *fog-2*; GFP::SP12 oocytes and detected two major types of CAR-1 condensates (Figure 4E). Some CAR-1 condensates appeared hollow, and the ER was detected on the outer surface of the condensates (Figure 4Ei), while other CAR-1 condensates were clearly not hollow but still partially co-localized with ER (Figure 4Eii, iii). Thus, both ectopic RNA-binding protein condensates in maturing oocytes and large RNA-binding protein condensates induced by extended meiotic arrest during development closely associate with ER sheets. To try to distinguish between the two possibilities of regulation, we asked if we could detect ectopic RNA-binding protein condensates without ER sheets or the converse. We generated a transient strain expressing GFP-tagged ER and mCherry-tagged MEX-3 and depleted *cct*. We observed multiple worms with very weak MEX-3 condensate phenotypes, but strong ER sheet phenotypes (Figure 4F). Since the ER sheets can clearly form in the absence of MEX-3 condensates, we conclude that expanded ER sheets may promote the condensation of RNA-binding proteins.

### De-repression of maternal mRNA translation correlates with ectopic condensation of regulatory proteins

Several of the RNA-binding proteins that become ectopically condensed after depletion of *cct* regulate the translation of maternal mRNAs in the *C. elegans* germline. CAR-1 and LIN-41 repress the translation of *spn-4*, while OMA-1 opposes LIN-41 to promote *spn-4* translation (Hubstenberger et al., 2013; Tsukamoto et al., 2017). To determine if there are disruptions in the function of the RNA-binding proteins when they are ectopically condensed, we depleted *cct-2* in a GFP::SPN-4::3xFLAG strain (Tsukamoto et al., 2017). In control oocytes SPN-4 was detected at high levels in the -1 oocyte as expected, and occasionally in the -2 oocyte (Figure 5A). In contrast, *cct-2* depletion resulted in high levels of SPN-4 in many oocytes, with significantly increased GFP::SPN-4 levels in the first five oocytes compared to the control (Figure 5B). In addition to being translationally derepressed, SPN-4 was ectopically condensed in most oocytes (Figure 5A). We next depleted *act-4* and observed an empty gonad in the proximal region and cellularized oocytes at the loop (Figure 5A; increased fluorescence levels in GFP-only image shown to allow clear visualization of cells at the loop). High levels of SPN-4 were detected in 2-4 oocytes near the loop in 73% of gonads, n= 15 (Figure 5A). We conclude that ectopic condensation of LIN-41 and CAR-1 may contribute to the translational de-repression of *spn-4* mRNA when *cct-2* is depleted. The phenotype after *act-4* depletion is harder to interpret because it was not as strong, and the severe disorganization of the gonad complicates the context. Nonetheless, the correlation between ectopic condensation and unregulated translation of *spn-4* occurs with both depletions.

**Figure 5.**
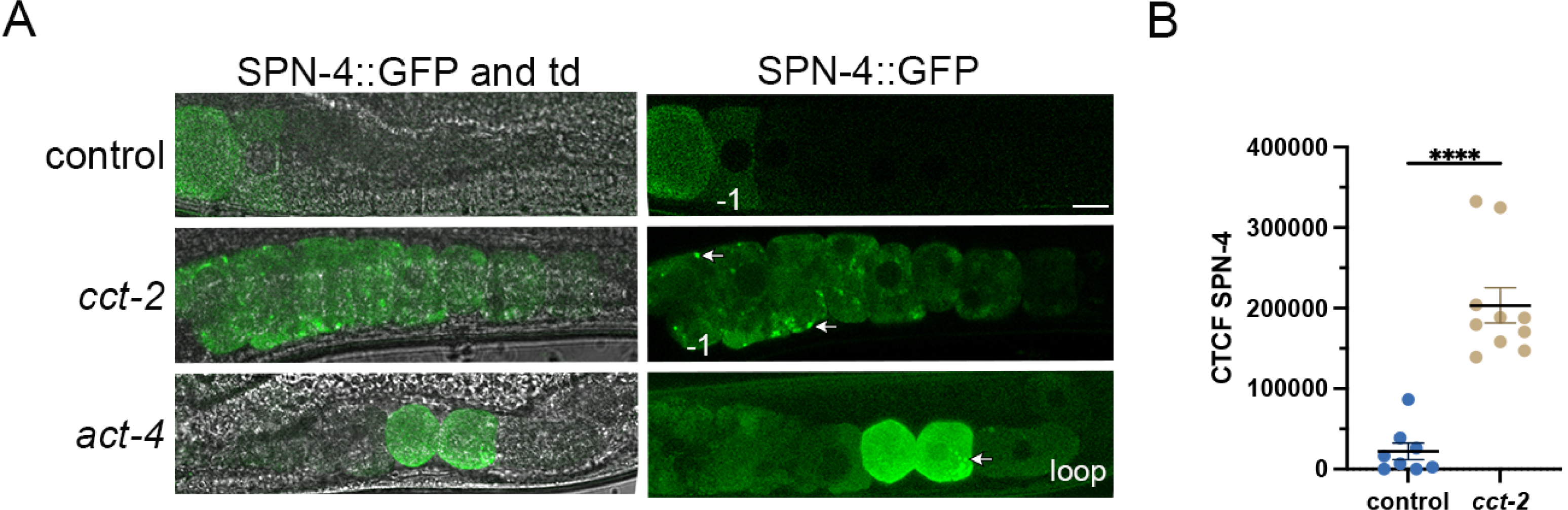
De-repression of maternal mRNA translation correlates with ectopic condensation of regulatory proteins. A) Confocal slices of GFP::SPN-4 in oocytes after RNAi of *cct-2* or *act-4* compared to the control *lacZ* RNAi. Arrows indicate ectopic clusters of SPN-4. Cellularized oocytes at the most proximal end are labeled -1 in control and *cct-2*, or near the loop in *act-4*. *Left)* Images of SPN-4::GFP, all at same levels and overlaid with transmitted light (td). *Right)* Images of SPN-4::GFP at same level for control and *cct-2,* and increased level for *act-4* to better see cells at the loop. B) Corrected total cellular fluorescence (CTCF) of SPN-4 levels in an ROI corresponding to five most-proximal oocytes. In the graph, error bars are mean +/-SEM. \*\*\*\**p*<0.0001. Scale bar is 10 µm in all micrographs.

## Discussion

Here we demonstrate the liquid-liquid phase separation of the conserved KH-domain MEX-3 protein *in vitro* and the complex regulation of MEX-3 condensation during oogenesis *in vivo.* Our findings strongly support that MEX-3 has intrinsic gel-like properties. We set out to investigate the extent to which the CCT chaperonin is required to maintain a decondensed phase of MEX-3 in maturing *C. elegans* oocytes. We identified a role for the folding activity of the CCT chaperonin oligomer in preventing ectopic condensation of select RNA-binding proteins in maturing oocytes. The CCT chaperonin modulates condensation of three regulators of maternal mRNA translation, CAR-1, LIN-41, and OMA-1, in addition to MEX-3. We also demonstrate a role for actin in this process, and our results suggest actin acts in parallel to and/or with the CCT chaperonin to regulate condensation of RNA-binding proteins. To further interrogate the regulation of condensation at a cellular level, we investigated the role of the endoplasmic reticulum. After depletions of *cct* subunits or actin, we detect dramatic remodeling of the ER into expanded sheets at the same time ectopic condensates form, as well as partial co-localization of RNA-binding protein condensates and the ER. Moreover, our partially penetrant RNAi conditions revealed that ER sheets can form before ectopic RNA-binding protein condensates, consistent with a model where expanded ER surfaces reduce the concentration threshold required for condensation. Lastly, we demonstrate ectopic condensation of regulatory proteins correlates with the de-repression of maternal mRNA translation, suggesting regulation of phase transitions is essential for RNA-binding protein function in oocytes.

### Properties of MEX-3 *in vitro* support *in vivo* phase separation

Several lines of evidence demonstrate MEX-3 can undergo liquid-liquid phase separation (LLPS) in physiologically relevant contexts. First, the increased condensation of recombinant MEX-3 in response to increased protein concentration and decreased salt concentration is consistent with MEX-3 undergoing LLPS (Alberti et al., 2019). Second, the spherical shape of the droplets indicates MEX-3 can condense. Third, our turbidity measurements in the presence of a crowding agent support our microscopy assays. Fourth, the observations that MEX-3 properties change over time, albeit on the scale of hours, support a phase-separated condensate (Patel et al., 2015; Alberti et al., 2018).

Our *in vitro* results provide further insights into the *in vivo* observations of MEX-3. In meiotically-arrested oocytes MEX-3 is detected in large granules that can be 10µm in diameter, but upon the resumption of meiosis, MEX-3 rapidly decondenses (Schisa et al., 2001; Elaswad and Watkins et al., 2022). MEX-3 associates with many other RNA-binding proteins and mRNAs in the large granules of arrested oocytes, and based on several functional assays has been thought to undergo phase separation *in vivo* (Elaswad et al., 2022). Mapping phase diagrams *in vivo* is challenging; however, our *in vitro* phase diagram demonstrates purified MEX-3 has the intrinsic ability to undergo LLPS. Our results strongly suggest MEX-3 is not liquid-like *in vitro*. The combination of no recovery in FRAP assays, resistance to hexanediol, absence of fusion events, resistance to elevated temperature, and stability over 30 mins all point towards either a gel-like or solid phase. We interpret the sensitivity of MEX-3 droplets to SDS and the changes in MEX-3 properties starting at 2 hours as being more consistent with a gel-like phase than a solid phase. Overall, the properties of purified MEX-3 are quite similar to MEG-3, a *C. elegans* germ granule protein that is gel-like (Wang et al., 2014; Putnam and Seydoux, 2019).

While preparing this manuscript, a study was published describing purified human Mex3A as undergoing LLPS in the presence of a crowding agent (Chen et al., 2024). Transcriptomic analyses indicate hMex3A is expressed at high levels in ovarian stromal cells and the spermatagonium (Le Borgne et al., 2014; Uhlén et al., 2015; Karlsson et al., 2021). The recombinant human Mex3A protein was investigated in three assays at concentrations ranging from 12.5-100 μM, significantly higher than the physiological range of 0.05-1 μM we used (Saha et al., 2016). hMex3A undergoes condensation and/or increased turbidity upon exposure to elevated temperature or increases in protein concentration in the presence of a crowding agent, similar to our findings for the *C. elegans* protein in the presence or absence of crowding agent. However, high salt induces condensation of 100 μM hMex3A protein, opposite of the effect we detected for *C. elegans* MEX-3. It is possible this difference is due to the differences in protein concentration used or the inclusion of a crowding agent. High salt concentrations can drive condensation in some heterotypic mixtures, as seen for FUS-A1 recently (Farag et al., 2023); however, the hMex3A study was not a heterotypic mixture. Despite this difference in results with hMex3 and MEX-3, the overall similar findings of LLPS are significant, especially considering the relatively low overall sequence identity between human Mex3A and *C. elegans* MEX-3 outside of the KH-domains (Buchet-Poyeau et al., 2007). We have previously speculated the phase separation of MEX-3 may be mediated by its intrinsically disordered regions (IDRs) which can promote weak multivalent interactions and are often found in phase-separating RNA-binding proteins (Wood et al., 2016; Alberti et al., 2019). This idea seems increasingly likely given the IDR regions of purified hMex3A are required for LLPS (Chen et al., 2024).

### The CCT chaperonin is a novel inhibitor of RNA-binding protein condensation during oogenesis

Our findings reveal a novel role for the CCT chaperonin during oogenesis as an inhibitor of RNA-binding protein condensation. While this function aligns with prior studies showing the CCT chaperonin inhibits P-body protein condensation and stress granules in yeast (Nadler-Holly et al., 2012; Jain et al., 2016), it was somewhat surprising because an opposing function had been identified in three prior genetic screens where CCT promotes condensation of RNA-binding proteins (Updike and Strome, 2009; Hubstenberger et al., 2013; Wood et al., 2016). Our study differed from those in that we focused on the regulation of RNA-binding protein phase transitions in actively maturing oocytes. We conclude that the folding activity of the CCT chaperonin oligomer is required to inhibit ectopic condensation of select RNA-binding proteins based on three lines of evidence: identical condensation phenotypes after depletion of each of seven *cct* subunits, decreased levels of β-tubulin after *cct* depletion, and highly mobile MEX-3 within clusters after FRAP experiments. The latter finding strongly suggests the MEX-3 clusters are not unfolded aggregates of MEX-3 protein because aggregates most often appear immobile, e.g. FRAP assays of amyloid and actin aggregates (Kim et al., 2002; Roberti et al., 2011; Saegusa et al., 2014). Therefore, this role of the CCT chaperonin is distinct from its role inhibiting and clearing aggregates of RHO-1 protein in *C. elegans* oocytes and suppressing monomeric mutant huntingtin (mHtt) aggregates in Huntington’s disease (Samaddar et al., 2021; Behrends et al., 2006). We tentatively conclude the CCT chaperonin is not required to inhibit ectopic condensation of PUF-3 or PUF-11 based on the absence of significant condensation after *cct* depletion. These results could be due to insufficient depletion during RNAi; however, the variable disorganization of oocytes caused by *cct* depletion in all experiments was observed. Importantly, the lack of ectopic condensation in the PUF-3/-11 oocytes supports the idea that effects on condensation are at least somewhat independent of oocyte organization in the gonad. At the same time, we acknowledge the pleiotropy of *cct* is a limitation in probing the mechanism of the CCT chaperonin in modulating condensation of RNA-binding proteins. Lastly, our *in vitro* condensation assays and prior studies of MEX-3 in arrested oocytes provide additional lines of evidence to support the conclusion that MEX-3 undergoes LLPS *in vivo* (Elaswad and Watkins et al., 2022). Therefore, we conclude RNA-binding protein phase transitions are most likely regulated by the action of the CCT chaperonin on one or more substrates in maturing oocytes.

### Actin and the ER may mediate part of the CCT chaperonin regulation of RNA-binding protein condensation

In considering how the CCT chaperonin might act to inhibit ectopic condensation of RNA-binding proteins, we first asked if *cct* depletion results in elevated levels of MEX-3. Our findings of a 43% increase suggest this may in fact be a contributing factor of the ectopic condensates. Because actin monomers are some of the best-characterized substrates of the CCT chaperonin, we also explored the possibility that actin mediates the *cct* phenotype (Frydman et al., 1992; Gao et al., 1992). Our first prediction if this model was correct would be very similar condensation phenotypes after depletion of *actin* and *cct*. Our analysis after RNAi of *actin* is complicated by the severe effects on germline organization, especially in the proximal region. Nonetheless, after depletion of *actin* we detected penetrant, ectopic condensation of MEX-3, CAR-1, and OMA-1, and weakly penetrant ectopic LIN-41 condensates. These condensates were similar to *cct* condensates in that they were enriched at the cortex and nuclear membrane of cellularized oocytes. MEX-3 condensates were also similarly mobile after FRAP. On the other hand, we also predicted that depletion of the obligate co-chaperone for CCT folding of actin, *prefoldin,* would result in similar condensation phenotypes, and none were detected. We are cautious interpreting these negative results with RNAi, especially because our controls examining the effect of *pfd* depletion on levels and polymerization of beta-tubulin indicated depletions were incomplete. Taken together, our results support a model where CCT functions in parallel to and/or with actin to regulate phase transitions of RNA-binding proteins (Figure 6). Our findings are the first to our knowledge to implicate actin as a critical regulator of RNA-binding protein condensation during oogenesis. In general, actin has not been implicated as a major regulator of cytoplasmic LLPS; however, branched actin networks promote condensation of p62 bodies in cell culture, and the actin nucleation factor N-WASP (Neural Wiskott-Aldrich syndrome protein) inhibits condensation of alpha-synuclein in muscle cells of *C. elegans* (Feng et al, 2022; Jackson et al., 2024). Human alpha-synuclein undergoes LLPS in neurons and its misfolding/ aggregation is associated with neurodegenerative diseases. Thus, we predict future studies may uncover additional more roles for actin regulating LLPS in diverse cell types.

**Figure 6.**
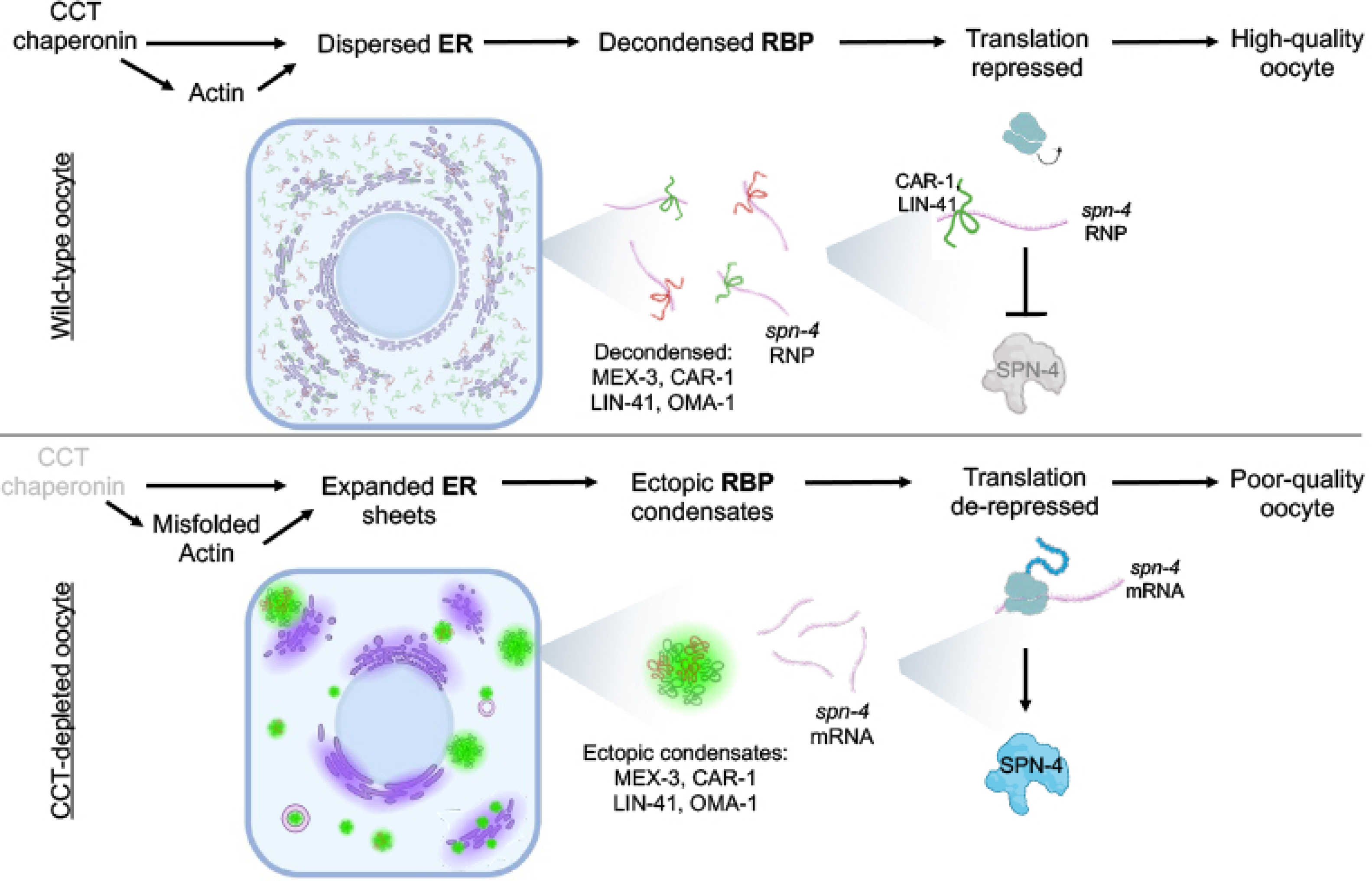
Model of RNA-binding protein phase transitions linking the CCT chaperonin, actin, and the ER with RNA-binding protein phase transitions in maturing *C. elegans* oocytes to regulate gene expression and promote oocyte quality. In maturing oocytes the CCT chaperonin and actin act coordinately and/or in parallel to promote a relatively dispersed ER distribution and decondensed phases of many RNA-binding proteins, some of which function to repress translation of maternal mRNAs like *spn-4* which is essential for high-quality oocytes. When the activity of the CCT chaperonin is depleted, actin is misfolded, and expanded sheets of ER result that are closely associated with ectopic condensates of RNA-binding proteins. In oocytes with ectopic condensates, translation of the LIN-41 and CAR-1 target mRNA *spn-4* is de-repressed which leads to poor-quality oocytes.

To further explore how cellular alterations might contribute to ectopic condensation phenotypes, and because membrane surfaces have emerged as essential regulators of condensation, we focused on the ER (Snead and Gladfelter, 2019; Snead et al., 2022). Prior studies revealed intriguing correlations between large RNA-binding protein condensates and ER sheets in *C. elegans* oocytes arrested in prophase an extended time and speculated the ER might recruit RNA-binding proteins into condensates (Patterson et al., 2011; Langerak et al., 2017). Here, we build on this model by demonstrating that expansion of ER sheets occurs concomitantly as ectopic RNA-binding protein condensation after depletion of *cct* and *actin*. Our data further suggest that the expanded ER sheets after *cct* depletion do not contain nuclear pore complexes, i.e. are not annulate lamellae. We uncovered a close association and partial co-localization between ER sheets and ectopic RNA-binding protein condensates. In some *cct-*depleted oocytes, the ER appears to wrap around a spherical condensate or reside directly adjacent to it, while other large, irregularly-shaped condensates appear to coat the ER sheets. In arrested oocytes, we detected some hollow RNA-binding protein condensates that appear to be surrounded by the ER. This striking association was somewhat reminiscent of the association between ADAD2-RNF granules and the ER in the mouse testis, where the ER appears to be located internal to the RNA-binding protein which is opposite of our observation in arrested worm oocytes (Chukrallah et al., 2023). *In vitro* systems have generated complex architectures where condensates can either be engulfed by membranes, or they can surround membranes, depending on several factors (Mangiarotti et al., 2023).

The partial co-localization between the ER and RNA-binding protein condensates after *cct* depletion and in arrested oocytes suggests the possibility of co-regulation. For example, studies with giant unilamellar vesicles describe membrane wetting and remodeling by condensates (Mangiarotti et al., 2023). In contrast, synthetic membranes promote the condensation of the *Ashbya gossypii* Whi3 RNA-binding protein at low protein concentrations (Snead et al., 2022). Thus, ectopic RNA-binding protein condensates may modulate the ER architecture in our system, or the increased ER surfaces in *C. elegans* oocytes could promote the condensation of RNA-binding proteins. The observation that depletion of CAR-1/Trailerhitch disrupts ER organization in early embryos/ nurse cells and oocytes could be seen to bolster the first model (Wilhelm et al., 2005; Squirrel et al., 2006). However, this model seems to be excluded by our observations of expanded ER sheets in the absence of ectopic RNA-binding protein condensates after weak depletion of *cct*. The idea of the ER contacting RNA-binding proteins to regulate their condensation in maturing oocytes is not particularly surprising. The ER physically connects to nearly every membrane-bound organelle and more recently is shown to be tethered with and regulate the numbers of P-bodies and stress granules in U2-OS cells (Lee et al., 2020). We are also struck by the possibility that in arrested *C. elegans* oocytes, the ER plays a role analogous to the mitochondria in prophase mammalian oocytes, where physical and functional interactions occur between mitochondria and the MARDO (mitochondria-associated ribonucleoprotein domain) (Cheng et al 2022). The MARDO contains several RNA-binding proteins including Lsm14/ CAR-1 and Ddx6/CGH-1, as well as vertebrate-specific RNA-binding proteins, and has roles in mRNA storage, translation, and degradation, but limited association with the ER. CAR-1 and CGH-1 are also condensed in the large RNP granules of arrested *C. elegans* oocytes (Jud et al., 2008; Noble et al., 2008; Elaswad et al., 2022). Thus, the cellular mechanisms that modulate phase transitions during oogenesis appear to be functionally conserved.

### Regulation of RNA-binding protein phase transitions may be essential to regulate maternal mRNAs

Protein and RNA function is dependent on its phase; for example, the liquid-to-solid phase transition of *oskar* RNP granules in *Drosophila* oogenesis is essential for regulation of localization, translation of Osk protein, and development (Bose et al., 2022). The ectopic condensation of multiple RNA-binding proteins that regulate translation of maternal mRNAs after *cct* and *actin* depletion provided an opportunity to ask if RNA-binding protein function was disrupted in these oocytes. We detected significantly increased levels of SPN-4 protein in expanded numbers of oocytes after depletion of *cct* and mildly increased levels after depletion of *actin*. Translation of *spn-4* mRNA is normally repressed by LIN-41 and CAR-1 in most growing oocytes and is promoted by OMA-1 (Hubstenberger et al., 2013; Tsukamoto et al., 2017). Our results suggest that LIN-41 and CAR-1 function may be disrupted because of ectopic condensation (Figure 6). In mammalian oocytes, the regulated phase separation of MARDO components is critical to both regulate mRNA translation and maintain oocyte quality (Cheng et al., 2022). Because increased levels of SPN-4 during oogenesis can interfere with progress to the diakinesis stage of prophase and contribute to infertility (Tsukamoto et a., 2017), the regulation of oogenic RNA-binding protein phase transitions may be critically important to maintain fertility. Collectively, our results provide important insights into how post-transcriptional modulation of the oocyte proteome is regulated.

In summary, our identification of the CCT chaperonin and actin as modulators of RNA-binding protein phase transitions in maturing oocytes provides insight into understanding a novel regulatory network that may be essential in the oocyte-to-embryo (OET) transition. The regulation of vertebrate maternal mRNA phase transitions during the OET was recently demonstrated in *Xenopus* and zebrafish, further underscoring the large-scale remodeling of maternal products that remains to be fully understood (Hwang et al., 2023). Beyond the implications of our findings for oocyte quality and fertility, hMex3A has been associated with multiple types of cancer (Mougeot et al.,2011; Jiang et al., 2012; Chao et al., 2019). In particular, hMex3A associates with P-bodies and circular RNAs to promote P-body dynamics and mRNA degradation in colorectal cells, leading to poor survival outcomes for patients (Chen et al., 2024). Thus, our findings for the conserved MEX-3 protein may also be applicable to better understanding the role of hMex3 phase transitions in cancer.

## Methods

### Worm strains and culture

*C. elegans* strains were cultured using standard protocols (Brenner, 1974) at 20C or 24C (for strains with GFP-tagged transgenes). The following strains were used, some of which were acquired from the CGC (Sternberg et al., 2024):

CB4108 *fog-2(q71)* V

DG4269 *mex-3(tn1753[gfp::3xflag::mex3])* III

DG3913 *lin-41(tn1541[GFP::tev::s::lin-41])* I,

DG4611 *puf-11(tn1824[puf-11::gfp::tev::3xflag])* IV

DG4607 *puf-3(tn1820[puf-3::gfp::tev::3xflag])* IV

DG4158 *spn-4(tn1699[spn-4::gfp::3xflag])* V

JH2054 *unc-119(ed3)* III; *axIs1492[pie-1p::GFP::tbb-2(orf)::tbb-2 3’ UTR + unc-119(+)]*

JH2184 *unc-119(ed3)* III; *axIs1595[pie-1p::GFP::npp-9(orf)::npp-9 3’UTR + unc-119(+)]*

OD61 *unc-119(ed3)* III; *ItIs41*[*pAA5; pie-1::GFP-TEV-STag::CAR-1; unc-119(+)]*

PHX8951 *mex-3a(syb8951[mCherry::mex-3])* III RT130 pwIs23*[vit-2::GFP]*

TX189 *unc-119(ed3)* III*; teIs1[(pRL475) oma-1p::oma-1::GFP + (pDPMM016) unc-119(+)]*

WH327 *unc-119(ed3)* III; *ojIs23[pie-1p::GFP::C34B2.10]*

DCL569 mkcSi13 II; *rde-1(mkc36)* V

Crosses were conducted using standard techniques to generate *fog-2*; GFP::SP12. Strains were synchronized before experiments using a hypochlorite bleaching method. In experiments with *fog-2* females, females were acquired as described and used two days after the L4 larval stage (Patterson et al., 2011).

### CRIPSR genome editing

Our first attempt to tag CCT-6 at the N-terminus with GFP resulted in larval lethality; therefore, we used the smaller 3xFLAG tag and followed the strategy of Zang et al (2018), targeting an insertion location that is unstructured at P373/K374 of CCT-6. The translational 3xFLAG-protein gene fusion to the endogenous *cct-6* locus was constructed using the method of Paix et al. (2017). The gRNA was designed using crispr.tefor.net and the IDT web-based tool: *cct-6* crRNA sequence: aucugugacucuucuuauca (IDT); *cct-6* repair template: agaaatacacgtttatcgaggaatgccgtgctccagactacaaagaccatgacggtgattataaagatcatgatatcgattacaaggatgacgatgacaagaagccactcacactactaatcaagggaccaaacaagcataccatcactcaaatc. Co-conversion reagents for *dpy-10* were used, resulting in a roller phenotype for screening; *dpy-10* crRNA: gcuaccauaggcaccacgag and repair oligonucleotide: acttgaacttcaatacggcaagatgagaatgactggaaaccgtaccgcatgcggtgcctatggtagcggagcttcacatg gcttcagacca acagcctat (Arribere et al., 2014; Paix et al., 2017). The edited *cct-6* locus was validated by PCR genotyping: forward primer: tcttgctcttcgtcgtgcc; reverse primer: gaacagcctctgaaaataggttact, and sequencing. We obtained two viable lines, one of which had low brood sizes.

### MEX-3 protein purification and labeling

MEX-3A full-length fused to an N-terminal 6XHis tag in a pTrcHis vector was expressed and purified from *E. coli* inclusion bodies under denaturing conditions following established protocols (Putnam and Seydoux, 2021). His-tagged MEX-3 was purified from solubilized inclusion bodies using an ӒKTA pure fast protein liquid chromatography (FPLC) system equipped with a 5 mL HisTrap HP nickel affinity chromatography column (Cytiva HisTrap HP). Fractions containing pure MEX-3 were identified via SDS-PAGE and Western blot and pooled together. MEX-3 was refolded into its native conformation via multistep dialysis in buffers containing decreasing concentrations of urea, imidazole and glycerol using established protocols (Putnam and Seydoux, 2021). The purity of refolded MEX-3 was assessed by SDS-PAGE and verified by Western blot using anti-His-tag antibodies (Supplementary Figure 1A).

MEX-3 was fluorescently labeled with DyLight-488 NHS Ester (ThermoFisher) (Putnam and Seydoux, 2021). To remove free dye, the labeling reaction was passed through Zeba™ Spin Desalting Columns (7K MWCO, 0.5mL; ThermoFisher). Fluorescent protein was verified by SDS-PAGE and a Typhoon gel imaging system. Trace-labeled MEX-3 solutions were prepared by mixing 488-labeled protein with unlabeled protein in a ratio of 1/10 labeled/unlabeled.

### In vitro Condensation Assays

Protein condensation was induced by diluting trace-labeled MEX-3 into condensation buffer containing 20 mM HEPES (pH 7.4) and salt adjusted to the desired final concentration. To complete the phase diagram, the final concentrations of trace-labeled MEX-3 were 0.05 μM, 0.1 μM, 0.5 μM, and 1 μM and salt adjusted to a final concentration of 50-1000 mM. Physiological concentrations of MEX-3 in oocytes (0.1 μM MEX-3) are based on the estimated MEX-3 concentration in embryos of 183 nM, (Saha et al., 2016). Physiological salt concentrations are 150 mM (Putnam and Seydoux, 2021). For crowding agent (PEG8000) experiments, protein was diluted in condensation buffer containing 3%, 5%, 10%, 15%, or 30% PEG8000.

For the turbidity assay, unlabeled MEX-3 was diluted in condensation buffer to 1 μM, and PEG8000 was used at 5%, 10%, 15%, or 30%. Absorbance measurements at 600 nm were obtained using a Nanodrop. Three measurements for each dilution were obtained in two independent experiments. Turbidity measurements were unreliable in the absence of crowding reagent and at low MEX-3 concentration (0.1 µM), likely due to the minimal condensation. For the time-course experiment, trace-labeled MEX-3 was diluted in condensation buffer to 0.1 μM and 150 mM salt, incubated at room temperature, and imaged at 0 min, 15 min, 30 min, 1 hr, 2 hr, 4 hr, 8 hr, and 24 hr timepoints.

For the heat exposure assay, trace-labeled MEX-3 was used at 0.1 μM in 150 mM salt. The tube was placed in a small volume of water equilibrated to 34°C in a 5 cm petri dish. The petri dish was kept in the Tokai Hit stage-top incubator pre-equilibrated to 34°C for the duration of imaging. At each timepoint: 0 min, 15 min, 30 min and 60 min, imaged immediately. The control was performed in parallel at room temperature and imaged at matching timepoints. For hexanediol (HD) and SDS experiments, trace-labeled MEX-3 was used at 0.1 μM or 1 μM, in buffer with 150 mM salt and 10% PEG. 10 μl of each condensation reaction was imaged to ensure condensates before adding 10% HD or 10% SDS.

To image all condensation assays, a Nikon A1R laser scanning confocal microscope was used with the 60x objective. Imaging was done within 10 mins of adding protein into condensation buffer unless noted. For each slide, 10 µm Z stacks (1 µm step size) from 5 random fields of views (214.24 x 214.24 µm) were acquired using automated XY image acquisition. A minimum of two independent replicates were done for each experiment. HV levels were adjusted for each MEX-3 concentration at 1M salt (where the protein was diffuse in the field of view) for optimal visualization of droplets. The same levels were used for all salt dilutions for a given protein concentration.

### RNAi

RNAi clones were obtained from the Source Bioscience RNAi library (Kamath and Ahringer, 2003), the Vidal library, or generated by PCR amplification and cloning of genomic sequences into the pL4440 vector. A vector with the bacterial *lacZ* gene was used as the negative control in all experiments. All gene identities in RNAi clones were verified by sequencing. We note the clone labeled *cct-4* in the Source Bioscience RNAi library is misidentified. Each colony of bacteria was grown in LB with 50 μg/ml carbenicillin at 37°C for 5-7 hrs and induced with 5 mM IPTG for 45-60 min at 37°C. Cultures were plated on RNAi plates (NGM plates containing 50 μg/ml carbenicillin and 1 mM IPTG). RNAi was performed by feeding L4-stage hermaphrodites for either 24 h at 24°C or 35-40 h at 20°C (*cct* genes). When possible, RNAi plates were blinded before image collection and analysis. A minimum of three replicates was performed for each RNAi experiment.

### Immunoblotting

Recombinant MEX-3 protein was mixed with 2x Laemmli in a 1:1 ratio and boiled at 95°C for 10 min before loading. For each RNAi-group, total protein was extracted from 50-100 adult hermaphrodites, following the protocol of Wang et al., (2016). Worms were picked into cold 1x PBS. Worm pellets were washed with 1x PBS to remove OP50 bacteria and resuspended in sample buffer containing 2x Laemmli, 3.5% SDS, 150 mM DTT and boiled at 95°C for 15 min before loading on gel.

Proteins were resolved on stain-free 12% Mini-PROTEAN polyacrylamide gels (Bio-Rad) and transferred to PVDF membranes activated in cold methanol. The blots were imaged on the Bio-Rad Chemidoc imaging system (stain-free blot setting) after transfer to ensure successful protein transfer. Blots were blocked for 30 min at room temperature in Tris-buffered-saline (TBS) with 0.05% Tween 20 (TBST) and 5% non-fat dry milk (TBS). Membranes were incubated with primary antibodies diluted in TBS for 2 hours at room temperature, washed three times for 10 min in TBST, incubated with secondary antibodies diluted in TBS for 1 hour at room temperature. Membranes were washed three times for 10 min in TBST, and visualized with Pierce ECL Western Blotting Substrate (ThermoFisher) and the BioRad Chemidoc imager. Semi-quantitative Western blots were repeated in triplicate.

Primary antibodies used were HRP-conjugated anti-His (Invitrogen 46-1009; 1:2000) and mouse anti-FLAG (Sigma F1804; 1:1000). The secondary antibody used was HRP-conjugated goat anti-mouse IgG (Jackson Lab 115-035-003; 1:10,000).

### Immunostaining

The anti-CAR-1 antibody was a gift from Dr. Keith Blackwell. Adult worms were dissected in M9 buffer to extrude their gonads and flash-frozen on dry ice. For experiments requiring GFP to persist post-fixation, we developed a fixation method that preserves GFP expression, although it is not optimal for CAR-1. Gonads were dissected in 5% paraformaldehyde (60 mM PIPES, pH 6.7) for 5 min at room temperature, then freeze-cracked in cold methanol for 5 min and washed in Tris-tween two times for 5 min each. For the experiment in the *rde-1* strain, gonads were dissected, frozen on dry ice, and fixed in ice-cold methanol for 1 min before adding 3.7% formaldehyde for 30 min at room temperature. Washes were as above. Chicken α-CAR-1 was used at 1:125. Alexa Fluor goat anti-chicken, 488 or 568 (Molecular Probes) was used at 1:200. Primary antibody incubations were performed at 4°C overnight, and secondary antibody incubations were performed at room temperature for 2 hr protected from light. A minimum of three replicates was performed for each experiment.

### Image acquisition of live and fixed worms and FRAP

Worms were imaged using either the Nikon A1R laser scanning confocal microscope or the Zeiss AXIO imager M2, equipped with Apotome3. Worms were picked onto slides made with 2% agarose pads and immobilized using 6.25 mM levamisole and a cover slip. Live worms on each slide were imaged within 10 mins to avoid inadvertent stress using the 60x objective (Nikon) or 40x objective (Zeiss) (Elaswad and Munderloh et al., 2022). Z stack images were acquired using 0.5 µm step size (Nikon) or 0.3 µm (Zeiss Apotome system). All images of a given strain in an experiment were collected using identical levels and settings.

For FRAP experiments, worms were immobilized in 25 mM levamisole on 6% agarose pads to eliminate minor worm movements. The 488 laser was used to photo-bleach an area of 3.68 μm^2^ in a -2 oocyte. Proteins were imaged using a 500-ms exposure and 60x objective. Images were collected over a 300s recovery phase, every 8s. ImageJ was used to correct for slight movements during imaging (see Image Analysis). A minimum of nine worms were measured for each experimental condition.

### Image Analysis

Quantitative analysis of *in vitro* condensation assays was performed using Fiji/ImageJ2/2.14.0 following methods from Putnam and Seydoux (2021). Maximum intensity projections of Z stacks were generated at each field of view. The brightness and contrast of all images were uniformly adjusted using histograms of intensity distributions to optimize visualization of condensates. For *in vitro* experiments, analysis was performed on at least five fields of view from two or more independent trials. Particle analysis was conducted using Fiji to quantify the number, integrated density (the total intensity of each object), area (size), and circularity of droplets or *in vivo* condensates. Skewness analysis was performed to measure the uniformity of fluorescence signal in an ROI, where relatively higher skewness values indicate less uniform distribution of fluorescence, indicative of condensation. In each worm, skewness was measured for each of the -1 to -3 oocytes, excluding the nucleus, and the mean value was reported. To calculate the corrected total cell fluorescence (CTCF), the integrated density and area of the -1 to -5 oocytes, and the mean fluorescence of background, was measured. CTCF= integrated density-(area of selected ROI x mean fluorescence of background).

To correct for worm movements during *in vivo* FRAP, the Stacks-Shuffling-MultiStackReg plugin in ImageJ was used to align time series to an image at t=0. Image analysis was performed using FIJI/ImageJ. Raw fluorescence intensity values of a bleached ROI and a reference ROI were corrected by subtracting background. For *in vivo* FRAP, the reference ROI was an unbleached granule, and the background was a cytoplasmic ROI. For *in vitro* FRAP, the reference ROI was the whole field, and the background was a ROI of the same size as the bleached ROI. Intensity values were normalized according to FRAP (t) = [I_B_(t) - I_B_(t_0_)]/[I_R_(t) - I_R_(t_0_)], where I_B_ is background-corrected fluorescence intensity of bleached zone at time (t) or bleach time (t_0_), and I_R_ is background-corrected fluorescence intensity of reference zone (unbleached granule for *in vivo* and whole field for *in vitro*) at time (t) or bleach time (t_0_) (Hubstenberger et al., 2013; Kang et al., 2015). Fractional recovery of fluorescence intensity graphs were generated by plotting the normalized data post-photobleaching against time of the recovery phase.

In Western blot experiments, band density was quantified using Bio-Rad Image Lab software. The intensity of each FLAG band was normalized to the intensity of total loaded protein in the corresponding lane in the blot-after-transfer. The densitometry values of the experimental RNAi samples were adjusted relative to a scaled value of 1 for the *lacZ (RNAi)* control.

### Statistical analysis

G Power 3.1 power analysis was used to determine sample sizes of worm experiments. Three biological replicates were conducted for all experiments unless specified. Data are presented as mean +/-SEM unless otherwise indicated. Data were plotted and statistical analyses were performed on GraphPad Prism 10.2.0 and specific statistical tests are noted in figure legends. P values < 0.05 were considered statistically significant.

## Supporting information

Supplemental figures with legends

## Acknowledgements

We acknowledge the contributions of Elizabeth Breton and Andrea Montalbano for performing preliminary experiments. We thank David Greenstein for strains and helpful discussion; Ekaterina Voronina for protocols; Tim Schedl, Xantha Karp, Lisa Petrella, and Schisa lab members for helpful discussion; Benjamin Swarts for use of the FPLC; and Keith Blackwell for the CAR-1 antibody. We thank Alex DeMattei for assistance with analyses. Some strains were provided by the CGC, which is funded by NIH Office of Research Infrastructure Programs (P40 OD010440). This work was funded by an NIH grant 1R15GM147844-01 to J. Schisa. We also acknowledge the CMU Office of Research and Graduate Studies for summer funding for undergraduates C. Munderloh and C. Bright, and the CMU College of Science and Engineering for support of graduate students M. Elaswad and V. Tice.

