## Supplemental figures with legends for "The CCT chaperonin and actin modulate the ER and RNA-binding protein condensation during oogenesis to maintain translational repression of maternal mRNA and oocyte quality"

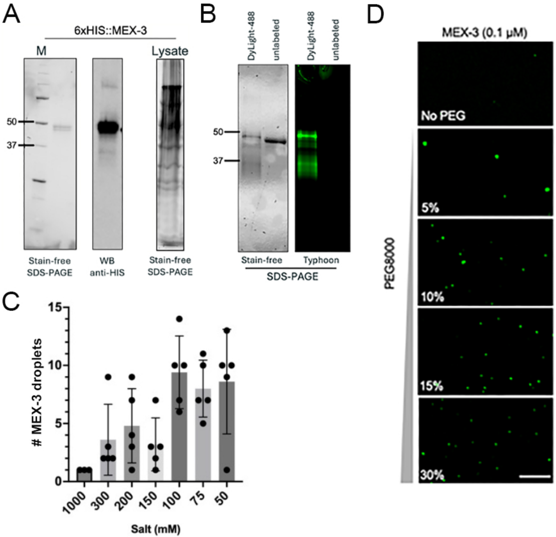

A
6xHIS::MEX-3
M
Lysate
50
37
Stain-free SDS-PAGE
WB anti-HIS
Stain-free SDS-PAGE
B
DyLight-488
untabeled
DyLight-488
untabeled
50
37
Stain-free
Typhoon
SDS-PAGE
C
### MEX-3 droplets
15
10
5
0
1000
300
200
150
100
75
50
Salt (mM)
D
MEX-3 (0.1 μM)
No PEG
PEG8000
5%
10%
15%
30%

**Supplementary Figure 1.** Fluorescently-labeled recombinant MEX-3 undergoes increased condensation under low-salt conditions and in the presence of a crowding agent. A) Purified MEX-3 on SDS-PAGE and anti-His Western blot. Multiple bands may correspond to isoforms of MEX-3. B) MEX-3 was fluorescently labeled using DyLight-488 and visualized on SDS-PAGE and using a Typhoon imager. C) The number of MEX-3 droplets increases as salt concentration decreases. D) Addition of PEG8000 to MEX-3 at 0.1 μM promotes droplet formation. Scale bar is 20 μm.

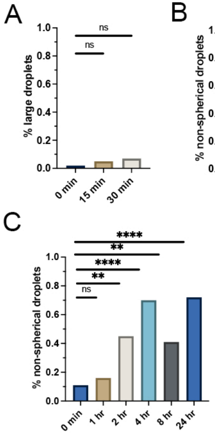
A
ns
ns
% large droplets
1.0
0.8
0.6
0.4
0.2
0.0
0 min
15 min
30 min
B
ns
ns
% non-spherical droplets
1.0
0.8
0.6
0.4
0.2
0.0
0 min
15 min
30 min
C
****
**
****
**
ns
% non-spherical droplets
1.0
0.8
0.6
0.4
0.2
0.0
0 min
1 hr
2 hr
4 hr
8 hr
24 hr

**Supplementary Figure 2.** MEX-3 does not appear liquid-like, but size and shape changes over time are consistent with a gel-like phase. A and B) No significant changes detected in the size or shape of droplets after 15 or 30 min at room temperature. C) The percent of non-spherical droplets, with circularity <0.8, increases significantly after 2 hours at room temperature and increases further at 24 hours. ns is not significant; * $p<.05$; ** $p<0.01$, **** $p<0.0001$ by Kruskal-Wallis test.

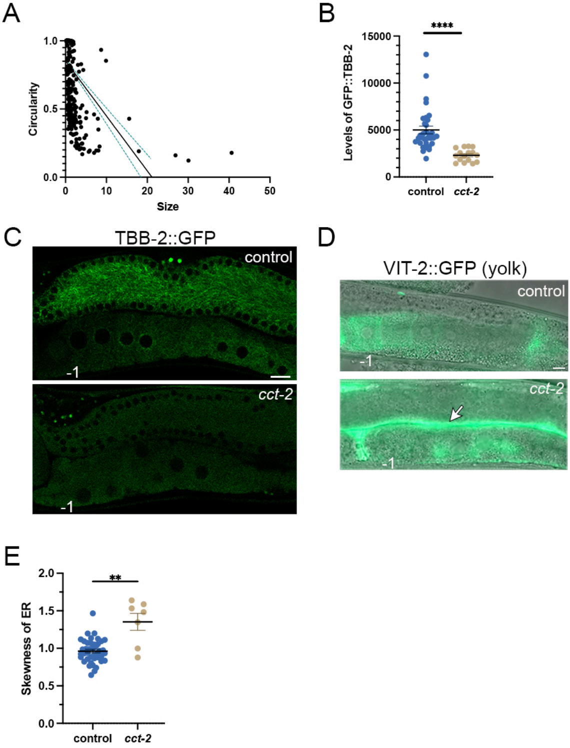

A
1.0
0.5
0.0
Circularity
0
10
20
30
40
50
Size
B
****
15000
10000
5000
0
Levels of GFP::TBB-2
control
cct-2
C
TBB-2::GFP
control
-1
cct-2
-1
D
VIT-2::GFP (yolk)
control
-1
cct-2
-1
E
**
2.0
1.5
1.0
0.5
0.0
Skewness of ER
control
cct-2

**Supplementary Figure 3.** Depletion of single *cct* subunits induces heterogeneous MEX-3 clusters, disrupts CCT chaperonin folding activity, and induces ER sheets. A) Circularity of MEX-3 clusters is negatively correlated with cluster size. $p<0.0001$ by Spearman test. B) Depletion of *cct-2* results in decreased GFP::TBB-2 (β-tubulin) levels in most proximal oocytes, indicating disrupted CCT chaperonin folding activity. GFP levels were measured in an ROI of the five most-proximal oocytes. C) Confocal single slices showing GFP::TBB-2 (β-tubulin) in germlines after depletion of *cct* subunit by RNAi compared to the control of *lacZ(RNAi)*. D) Fluorescence micrographs showing VIT-2:GFP as a marker of yolk in control oocytes and after *cct-2* RNAi. The -1 oocyte is marked, and the arrow points to large amounts of yolk that appear blocked from entering *cct* oocytes. E) Skewness assay shows significantly increased asymmetry of the ER in oocytes after depletion of *cct-2* compared to the control of *lacZ(RNAi)* (see Figure 4A). Scale bar is 10 μm in all micrographs.

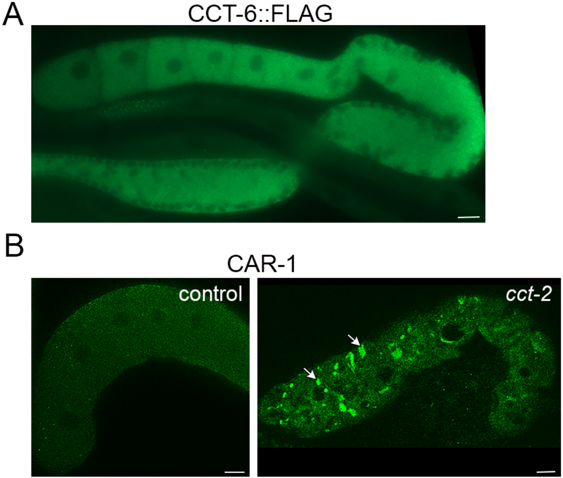
A
CCT-6::FLAG
B
CAR-1
control
cct-2

**Supplementary Figure 4.** The CCT chaperonin is expressed at high levels in the germline and acts cell autonomously. A) Fluorescence image of anti-FLAG immunostaining in CCT::FLAG germline. Note the disruption in staining of the distal gonad is due to the intestine atop the gonad.  B) Apotome image of anti-CAR-1 immunostaining in control and *cct-2(RNAi)* oocytes. Arrows indicate nuclear-associated and cortical condensates. Scale bar is 10 μm in all micrographs.

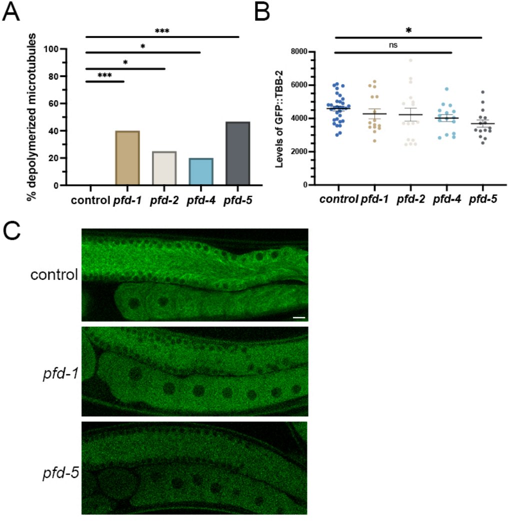

A
B
C
***
*
*
***
% depolymerized microtubules
100
80
60
40
20
0
control
pfd-1
pfd-2
pfd-4
pfd-5
*
ns
Levels of GFP::TBB-2
8000
6000
4000
2000
0
control
pfd-1
pfd-2
pfd-4
pfd-5
control
pfd-1
pfd-5

**Supplementary Figure 5.** Depletion of *pfd* subunits by RNAi is partially effective. A) The percent of gonads with depolymerized microtubules (GFP::TBB-2 used as β-tubulin reporter) is significantly increased after depletion of *pfd* subunits as compared to the *lacZ* control; however, the phenotype is <50% penetrant. B) Levels of GFP::TBB-2 after most *pfd* depletions were not significantly decreased. C) Confocal single slice images of GFP::TBB-2 gonads are representative images showing the decrease in microtubule polymerization seen in 20-47% of *pfd* depletions, and the modest decrease in GFP::TBB-2 levels after *pfd-5(RNAi)*. Scale bar is 10 μm. Error bars are mean +/- SEM. ns is not significant; * $p<.05$; ***$p<0.001$ by Kruskal-Wallis test.
